# The miR156/SPL9 interaction mode regulates the anthocyanin accumulation in potato

**DOI:** 10.1101/2025.01.06.631517

**Authors:** Nan Li, Hushuai Nie, Xiaojuan Wu, Peijie Wang, Yu Ma, Enze Yang, Juan Wu, Zhicheng Zhang, Rui Xie, Dan Wang, Yanhong Ma

## Abstract

Anthocyanins are an essential class of flavonoids that represent a large group of plant secondary metabolites. microRNAs (miRNAs) can target TFs related to anthocyanin synthesis and inhibit their expression, thereby affecting the expression level of key downstream structural genes and ultimately regulating the synthesis and accumulation of anthocyanins in plants. Nevertheless, the regulatory function of miR156a in anthocyanin synthesis, especially in potatoes, has yet to be investigated. In this study, the miR156a and its target gene *StSPL9* were screened and analyzed from small RNA sequencing, degradome sequencing and transient expression assay. The function of miR156a in anthocyanin synthesis of potato tuber was investigated. Overexpression of miR156a (OE-miR156a) in potato tubers promoted anthocyanin synthesis while concurrently reducing flavonoid synthesis compared with wild-type (WT) potatoes. The miR156a-silenced tubers (STTM-miR156a) achieved using short tandem target mimics (STTM) contained significantly lower anthocyanin content and increased flavonoid accumulation compared to WT and OE-miR156a. Notably, overexpression of *StSPL9* in potato yielded identical results. The relative expression levels of the anthocyanin-related structural genes *PAL*, *4CL*, *CHS*, *CHI*, *F3H*, *DFR*, and *UFGT* in STTM-miR156a transgenic potato tubers showed the opposite trend to that observed in OE-miR156a potato tubers, as demonstrated by RNA sequencing and quantitative real-time PCR (qRT-PCR). The integrated analysis of the transcriptome and metabolome revealed that the accumulation of flavonoids and flavonols in STTM-miR156a was significantly increased than in the wild type, whereas the contents of anthocyanins and phenolic acids were notably reduced. Further analysis showed that the expression of *CHS*, *FG2*, *BEATH*, and *CYP75A* were up-regulated in STTM-miR156a, and *HCT* was downregulated, which resulted in a reduced accumulation of methyl 4-caffeoylquinate, a phenolic acid metabolite. Therefore, it is demonstrated that miR156a indirectly modulates the expression of *CHS*, *BEATH*, *CYP75A*, *HCT*, and *FG2* by targeting and degrading its target gene *StSPL9* and other *SPL* genes, thus influencing anthocyanin accumulation in potato tuber. This study provides valuable insights into the complex regulatory network governing anthocyanin biosynthesis in *Solanum* species.

## Introduction

Potato (*Solanum tuberosum* L.) is the third most cultivated food crop globally. It provides essential nutrients, including carbohydrates, vitamins, and minerals, such as potassium, magnesium, and iron, which enhance the body’s immunity. Anthocyanins are phenolic water-soluble compounds in certain tropical fruits, red or purple-blue vegetables, grains, and tubers. Although anthocyanins are present in small amounts in the human diet, numerous studies have demonstrated their health advantages, suggesting they may help prevent various chronic diseases. Research indicates that anthocyanins can be effective in preventing metabolic disorders and microbial infections, including cardiovascular disease, cancer, and diabetes (Khoo et al, 2017) Additionally, they may improve conditions such as obesity dyslipidemia, hypertension, and blood glucose abnormalities(Amiot et al, 2016).

Color changes are readily observable during plant growth and development, particularly during the ripening of fruits and the senescence of leaves. These alterations in color significantly impact the nutritional and quality attributes of crops, especially those that produce tubers and fruits. Anthocyanin-rich colored potatoes play an important role in enhancing health and helping to prevent cardiovascular and cerebrovascular diseases. With the improvement of living standards, the nutritional demands for food are gradually increasing. Up to now, although some certain transcription regulators, plant hormones, and radiation have been shown to influence anthocyanin accumulation in potato tubers (Cui et al, 2023; Chen et al, 2024), our understanding of the regulatory mechanisms governing anthocyanin metabolism in colored potato tubers remains limited (Gonzalez et al, 2008; Van der Krol et al, 1990; Sun et al, 2019; D’Amelia et al, 2022). Since the potato is a homologous tetraploid crop capable of asexual reproduction and is sensitive to genetic homogeneity, it is essential to investigate the mechanisms regulating anthocyanin production through the genetic improvement of colored potato varieties. The biosynthetic genes and regulatory factors associated with anthocyanin production have been extensively studied in the context of fruit color change (Lu et al, 2017). However, the underlying principles governing pigment accumulation and the resulting color change are still largely unknown.

MicroRNAs (miRNAs) are a class of endogenous, non-coding small RNAs (19-23nt) that primarily regulate the expression of target genes at the post-transcriptional level, playing a crucial role in plant development (Luan et al, 2023). Among these, miR156 is one of the most highly conserved miRNA families in plants. Research has shown that the miR156/*SPL* module plays a vital role in plant freezing tolerance (Zhao et al, 2022), drought tolerance (Arshad et al, 2017), the accumulation of reactive oxygen species, and the salicylic acid signaling pathway (Yin et al, 2019). Furthermore, miR156 delays the vegetative phase transition of plants by controlling its target gene *SPL* (squamous promoter binding protein-like 9) (Zhang et al, 2019). It also influences leaf growth and number (Wang et al, 2019), inhibits tillering and branching (Zhang et al, 2021), affects fruit ripening (Manning et al, 2006), regulates the number of lateral roots (Yu et al, 2015), and impacts floral organ development, inflorescence development and plant fertility (Wang et al, 2016).

The miR156/*SPL* regulatory network governing anthocyanin accumulation has been recognized in multiple species, including *Arabidopsis,* for example (Mallory et al, 2004; Rhoades et al, 2002). overexpression of miR156 was found to decrease the expression of its target gene *SPL*, resulting in delayed flowering in the overexpressed plants. The miR156 targeted *SPL9* transcription factor competes with TT8 (Transparent testa 8) for binding to PAP1, thereby interfering with the MBW transcription complex and inhibiting the expression of *DFR*, which in turn suppresses anthocyanin synthesis (Gou et al, 2011). Additionally, Csn-miR156d regulates flowering and anthocyanin accumulation in tea plants by suppressing the expression of *CsSPL1* (Lin Q, 2024). In ‘Suri’ pear (*Pyrus pyrifolia Nakai*), the miR156/*SPL* module inhibits anthocyanin accumulation by inhibiting the expression of *PpMYB10* (Qian et al, 2017). In grapes, miR156/*SPL*s are involved in forming anthocyanins by responding to hormone signals such as ABA, MeJA, and GA (Su et al, 2021). In peach (*Prunus persica*), *PpSPL1* can reduce the expression level of *PpMYB10.1* (Zhou et al, 2015).

In addition, miR156 is also involved in the regulation of anthocyanin accumulation in *Arabidopsis*, blueberry and, other plants (Zhou et al, 2022; Gou et al, 2024; Wang et al, 2020; Li et al, 2024). Although the molecular mechanism involving miR156/*SPL* in the regulation of anthocyanin accumulation has been widely studied, the synthesis of anthocyanin regulated by miR156/*SPL* in potato tubers is not well studied. Consequently, a comprehensive investigation into the connection between miR156/SPL and anthocyanin is crucial for a deeper understanding of the intricate regulatory framework governing potato anthocyanin production. This study will also yield valuable insights for potential quality management.

In this study, we conducted a functional identification of miR156 and *SPL9* in potatoes. Our findings indicate that the miR156a and *StSPL9* genes regulate the color change of potato tubers. Notably, our research results revealed that the miR156a/*StSPL9* module serves as a crucial upstream regulatory factor, potentially influencing several essential regulatory genes in the anthocyanin and flavonoid pathways, including *CHS*, *BEATH*, *CYP75A*, *HCT*, and *FG2*. These results reveal a novel regulatory network mediated by the miR156a-*StSPL9* module, which plays a crucial role in the synchronized regulation of anthocyanin accumulation in potato tubers.

## Results

### In vitro validation of miR156a and its target gene *StSPL9*

In our previous research, we conducted small RNA and degradome sequencing on purple potato tubers during various developmental phases. Throughout the growth period, the expression level of miR156a showed an increase. Combined with the degradome sequencing, the target gene of miR156a was predicted to be *SPL9*, and the binding site was at the 1000-1500th base of the target gene (Figure 1a) (Wu et al, 2022). To further verify that miR156a targets StSPL9, the transient expression assay through *Agrobacterium tumefaciens* in tobacco leaves was performed. We injected pCAMBIA1302-mGFP5, OE-StSPL9-mGFP5, OE-miR156a-mGFP5 and the combination of OE-StSPL9+OE-miR156a-mGFP5 into the leaves of *Nicotiana Benthamiana* respectively (Figure. 1b), qRT-PCR detection of relative expression levels of mGFP5 and StSPL9.

**Figure 1.**
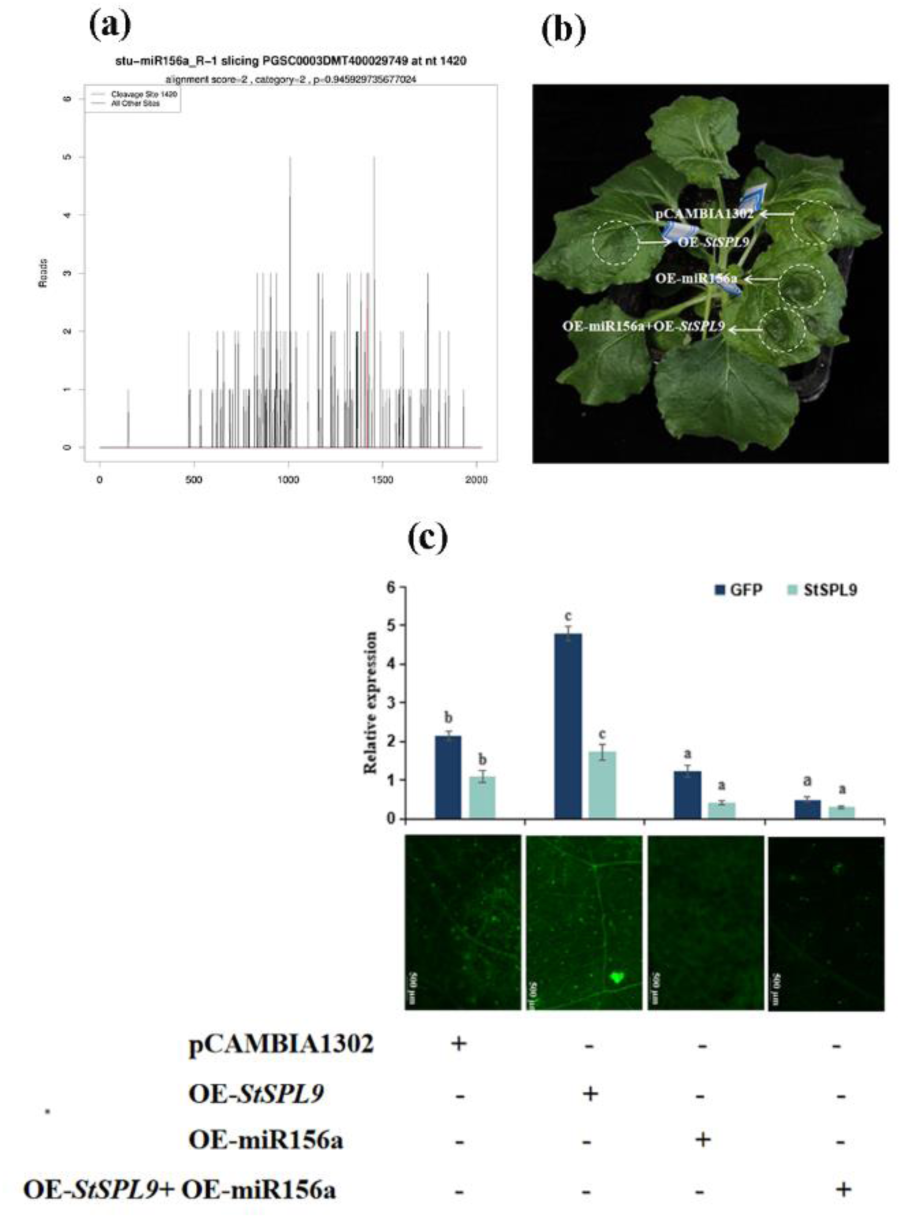
Verification of the interaction between miR156a and its target gene StSPL9. **a**: The binding site of miR156a to the target gene StSPL9 in the degradation group; **b**: pCAMBIA1302-mGFP5, OE-StSPL9-mGFP5, OE-miR156a-mGFP5 and OE-StSPL9+OE-miR156a-mGFP5 were injected into Nicotiana benthamiana leaves. **c**: miR156a qRT-PCR was performed on the leaves of Nicotiana benthamiana injected with miR156a and its target gene StSPL9; the luorescence intensity of Nicotiana benthamiana leaves injected with pCAMBIA1302-mGFP5, OE-StSPL9-mGFP5, OE-miR156a-mGFP5 and OE-StSPL9+OE-miR156a-mGFP5 was observed by confocal microscopy, and the relative expression levels of GFP and StSPL9 were corresponding.

The findings indicated that the relative expression trend of mGFP5 and StSPL9 in different leaves was consistent, and the expression level in OE-StSPL9-mGFP5 injection leaves was significantly higher than that in OE-StSPL9-mGFP5+OE-miR156a combination injection leaves (Figure. 1c). We utilized confocal microscopy to observe the fluorescence signal of mGFP5, revealing that the fluorescence intensity in tobacco leaves injected with OE-StSPL9-mGFP5 was the highest, while the intensity in leaves injected with OE-StSPL9-mGFP5+OE-miR156a-mGFP5 was the lowest (Figure. 1c). These results indicate that miR156a effectively recognizes and cleaves its binding sites within StSPL9, subsequently suppressing its expression.

### StSPL9 is localization of the nucleus

A recombinant vector called pCAMBIA1302-StSPL9-GFP, designed to encode the green fluorescent protein (GFP), was constructed to ascertain the subcellular localization of StSPL9.The cellular localization of StSPL9 was subsequently analyzed through transient expression in tobacco leaves. Results from confocal microscopy indicated that the control vector pCAMBIA1302-GFP was distributed throughout all regions of the tobacco epidermal cells. Conversely, pCAMBIA1302-StSPL9-GFP was specifically localized to the nucleus (Figure. 2), suggesting that StSPL9 may primarily function within the nucleus.

**Figure 2.**
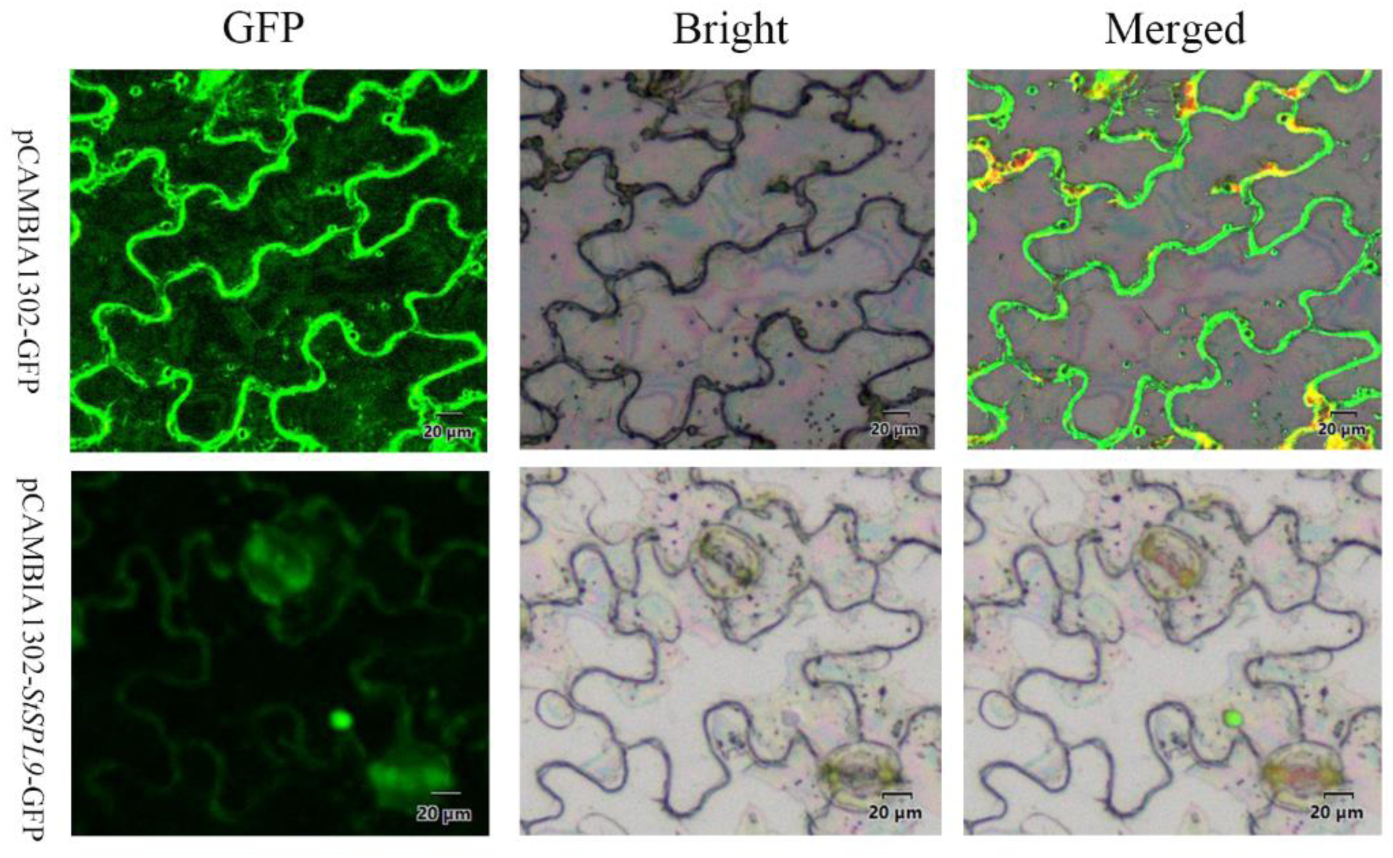
Subcellular localization of *StSPL9* in tobacco epidermal cells. The fusion protein pCAMBIA1302-StSPL9-GFP) and the GFP-positive control (pCAMBIA1302-GFP) were independently transiently expressed in tobacco cells. Scale bar=20 um.

### Regulation of anthocyanins and flavonoids accumulation in potato tubers by miR156a and *StSPL9*

To explore the role of miR156a, we constructed an overexpression vector that includes the precursor sequence of miR156a regulated by the 35S CaMV promoter, as well as an STTM vector control led by P35S (Figure S1a). Stem segments of potatoes in tissue culture were utilized for the transient silencing and overexpression of miR156a through *Agrobacterium-mediated* transformation, leading to the generation of 12 kanamycin-resistant and 13 hygromycin-resistant transgenic lines of miR156a. PCR identification was conducted using the STTM-35seq-F/NOSseq-R and pCAMBIA1302-miR156-F/R specific primers respectively (Table S1). In the STTM-miR156 transgenic line, an amplification product of approximately 238 bp was observed, while a product of about 423 bp was amplified in the OE-miR156a transgenic line. No amplification product was detected in the non-transgenic control line or the blank control (Figure S1b and c).

Wild-type, STTM-miR156a -10, 12, 13, and OE-miR156a -7, 10, 12 transgenic lines were cultivated in pots (Figure 3a). We extracted total RNA from the third leaves and tubers of wild-type, STTM-miR156a-10, 12, 13 and OE-miR156a -7, 10, 12, potato plants for qRT-PCR detection. The relative expression of miR156a in OE-miR156a and STTM-miR156a transgenic lines was found to be opposite. The levels of miR156a gene expression increased by 1.3 to 3.4 folds in leaves, while in tubers, an increase of 1.9 to 4.4 folds was observed (Figure 3b). As anticipated, in the STTM-miR156a transgenic lines, the expression of miR156a in the leaves of lines STTM-miR156a -10, 12, 13 was down-regulated by 0.45-0.62 folds when compared to the wild-type plants. In tubers, the expression of miR156a in three transgenic lines was down-regulated by 0.2-0.5 folds relative to the wild-type plants (Figure 3c). The *StSPL9* gene showed a relative expression 0.1-0.7 folds in OE-miR156a tubers. (Figure 3c), whereas in STTM-miR156a, the relative expression of the *StSPL9* gene was significantly increased by 1.4-3.6 folds (Figure 3b). These findings indicate that miR156a regulates the expression of StSPL9, confirming that it is a target of this microRNA.

**Figure 3.**
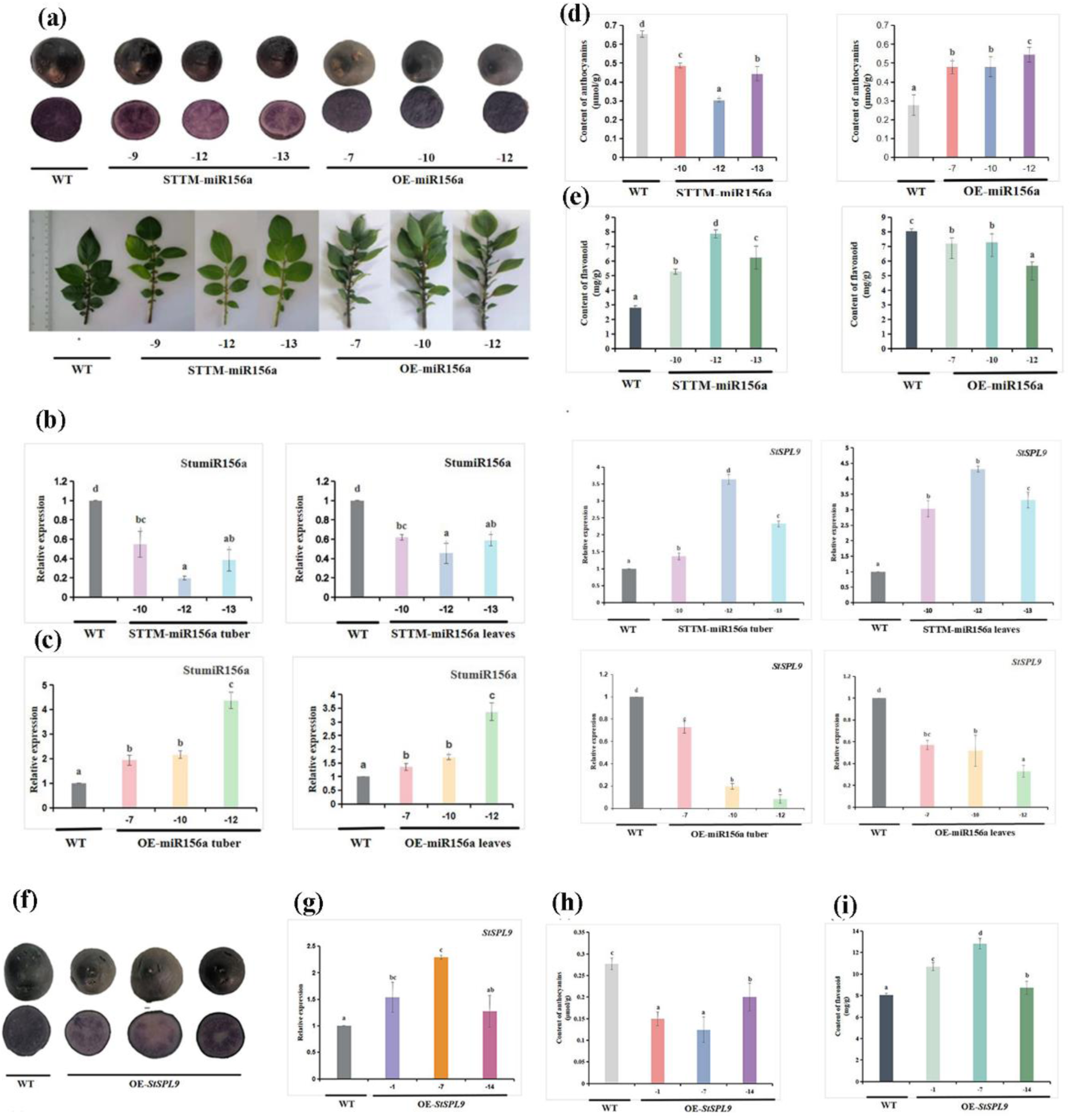
miR156a affected anthocyanin accumulation. **a:** Wild-type, potato tuber and leaf phenotypes after miR156a overexpression and silencing. The upper side represents cross sections of WT, STTM-miR156a (-9, -12, -13), and OE-miR156a (-7, -10, -12) potato tubers. Lower side represents WT, STTM-miR156a (-9, -12, -13), OE-miR156a (-7, -10, -12) potato pot leaves. **b:** The relative expression levels of miR156a and its target gene *StSPL9* in leaves and tubers of WT and STTM-miR156a transgenic lines were detected by qRT-PCR; **c:** The relative expression levels of miR156a and its target gene *StSPL9* in leaves and tubers of WT and OE-miR156a transgenic lines were detected by qRT-PCR. **d:** The anthocyanin content of WT and three STTM-miR156a and three OE-miR156a transgenic potato lines was determined; **e:** The contents of flavonoids in tubers of WT, 3 STTM-miR156a and 3 OE-miR156a transgenic potato lines were determined; **f:** Comparison of non-transgenic potato tubers (WT) and OE-*StSPL9* transgenic potato; **g:** The relative expression of *StSPL9* in OE-*StSPL9* transgenic potato tubers was verified by qRT-PCR; **h:** Determination of total anthocyanin content in OE-*StSPL9* transgenic potato tubers; **i:** Determination of total flavonoid content in OE-*StSPL9* transgenic potato tubers. The bar chart is the average±SD of three biological replicates. The letters in the same histogram indicate the significant difference between the values tested by the least significant difference (LSD), (p <0.05).

Notably, in contrast to the wild-type, the third leaf above the potted seedlings of STTM-miR156a transgenic lines exhibited a pronounced green coloration, while the tubers displayed a significantly lighter color. Conversely, the tubers of OE-miR156a transgenic lines exhibited a distinct purple hue (Figure 3a). Our findings indicated that the anthocyanin content in the tubers of STTM-miR156a transgenic potato were lower by a factor of 0.5 to 0.7 compared to the wild type (Figure 3d). In contrast, the tuber anthocyanin content in the OE-miR156a transgenic lines was up-regulated by 1.7 to 2.0 folds. Given that the anthocyanin biosynthesis pathway is generally part of the flavonoid pathway, we also assessed the flavonoid content. Compared to the wild type, the flavonoid content in STTM-miR156a transgenic potato tubers was significantly increased, showing an up-regulation of 1.9-2.8 folds. Conversely, in the OE-miR156a lines, the flavonoid content was significantly reduced and down-regulated by 0.7-0.9 folds (Figure 3e). These results indicate that miR156a plays a role in the regulation of anthocyanin and flavonoid accumulation in potato.

In our study of the role of *StSPL9* in potatoes, we established transgenic lines that overexpression *StSPL9* and selected three independent lines, designated OE-*StSPL9* -1, 7, 14. In the OE-*StSPL9* transgenic potato tubers, the relative expression of *StSPL9* was significantly up-regulated by 1.3-2.3 folds, and the tubers exhibited a noticeably lighter color compared to the wild type (Figure 3f). Concurrently, we measured the anthocyanin and flavonoid content in the *StSPL9* transgenic potato tubers. As illustrated in Figure 3h and i, the anthocyanin content in the three *OE-StSPL9* transgenic potato tubers was down-regulated by 0.4-0.7 folds, while the flavonoid content was significantly increased by 1.1-1.6 folds. The findings align with those from STTM-miR156a transgenic potatoes, indicate that *StSPL9* inhibits the biosynthesis of anthocyanins in potatoes, while simultaneously enhancing the synthesis of flavonoids.

### miR156a changed the expression levels of other genes in potato

Compared with wild-type, RNA-seq analysis of transgenic potato plants expressing STTM-miR156a and OE-miR156a identified 154 and 4,693 differentially expressed genes (DEGs), respectively. Of these, 103 and 2,458 DEGs were up-regulated in the STTM-miR156a and OE-miR156a transgenic potato plants, were then subjected 51 and 2,235 DEGs were downregulated (Figure 4a and 4c). The KEGG pathway of 154 and 4693 DEGs from STTM-miR156a vs WT and OE-miR156a vs WT were classified into 43 and 137 KEGG terms (Figure S2a and b). Most of the pathways were enriched in Metabolic pathways, Biosynthesis of secondary metabolites, Phenylpropanoid biosynthesis, Plant-pathogen interaction ((Figure 4b and d). Notably, Phenylpropanoid biosynthesis, Flavone and flavonol biosynthesis, and Flavonoid biosynthesis were more significantly enriched in the STTM-miR156a vs WT (Figure S2a and b).

**Figure 4.**
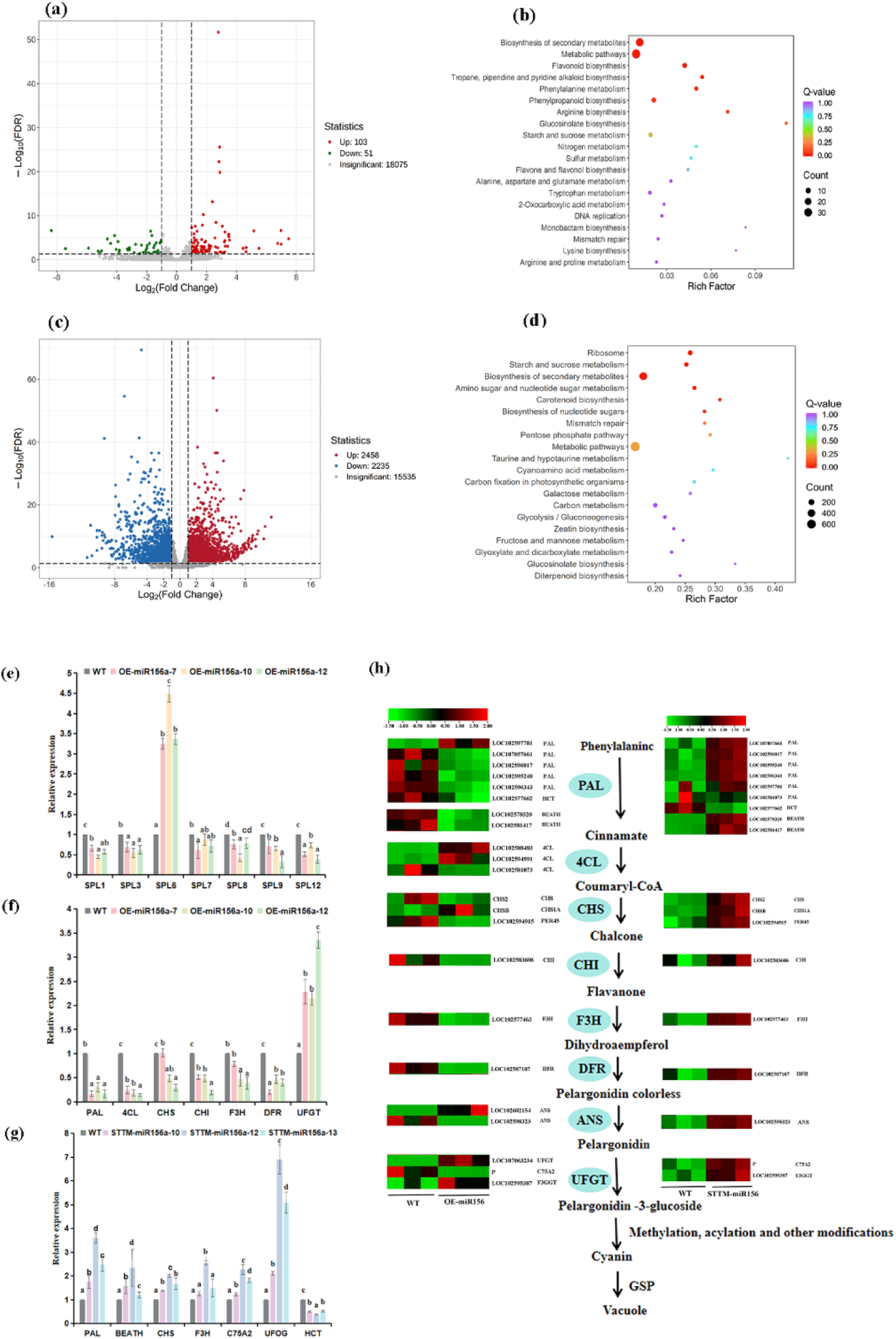
Transcriptomic analysis of wild-type and potato plants transgenic. **a:** The DEGs volcano map in STTM-miR156a. Note: The red dot represents the up-regulated differential gene, the green dot represents the down-regulated differential gene, and the gray band you represent the non-differentially expressed gene. **b:** KEGG enrichment scatter diagram of STTM-miR156a differential gene. Note: The abscissa represents Rich factor, the greater the Rich factor, the greater the degree of enrichment, and the ordinate represents the KEGG pathway; the larger the point, the more the number of differential genes enriched in the pathway; the redder the color of the point, the more significant the enrichment. **c:** DEGs volcano plot in OE-miR156a. **d:** KEGG enrichment scatter diagram of OE-miR156a differential gene. **e:** Relative expression levels of seven SPL genes screened from transcriptome data in OE-miR156a. **f:** Seven anthocyanin structural genes were screened in OE-miR156a for qRT-PCR to identify the accuracy of transcriptome information (*PAL*, *4CL*, *CHS*, *CHI*, *F3H*, *DFR*, and *UFGT*). **g:** The accuracy of transcriptome information was identified by qRT-PCR for 7 differentially expressed genes screened in STTM-miR156a, and the regulatory effect of miR156a on anthocyanin was further explained (*PAL*, *BEATH*, *CHS*, *F3H*, *C75A2*, *UFOG*, and *HCT*). **h:** Transcriptional profiles of anthocyanin biosynthesis genes in WT and OE-miR156a strains (left) and WT and OE-miR156a strains (right), heatmaps were generated based on transcriptome data. Statistical analysis was performed using one-way analysis of variance (ANOVA). Significant levels are denoted by asterisks: *P < 0.05.

The regulation of anthocyanin biosynthesis involves transcription factors, and we screened differentially expressed the transcription factors in STTM-miR156a and OE-miR156a. Both STTM-miR156a and OE-miR156a have *NAC*, *MYB*, *Trihelix*, *WRKY*, and *MADS*. It is noteworthy that in OE-miR156a also has *C3H*, *AUX/IAA*, *GARS*, *C2C2*, *bZIP*, *C2H2*, *bHLH*, *AP2*, and *SBP* (Figure S2d and e). *StSPL1*, *StSPL3*, *StSPL6*, *StSPL7*, *StSPL8*, *StSPL9* and *StSPL12* were differentially expressed from the RNA-seq of STTM-miR156a and OE-miR156a transgenic potato (Figure S3 and Table S3). Among them, the relative expression levels of *StSPL6* were significantly increased, and the relative expression levels of *StSPL1*, *StSPL3*, *StSPL7*, *StSPL8*, *StSPL9* and *StSPL12* were significantly decreased in OE-miR156a. Concurrently, qRT-PCR was employed to assess the expression levels of the seven *SPL* genes, and the findings aligned with those from RNA-seq. (Figure 5e). It is speculated that *SPL* genes were targeted and regulated by miR156a.

**Figure 5.**
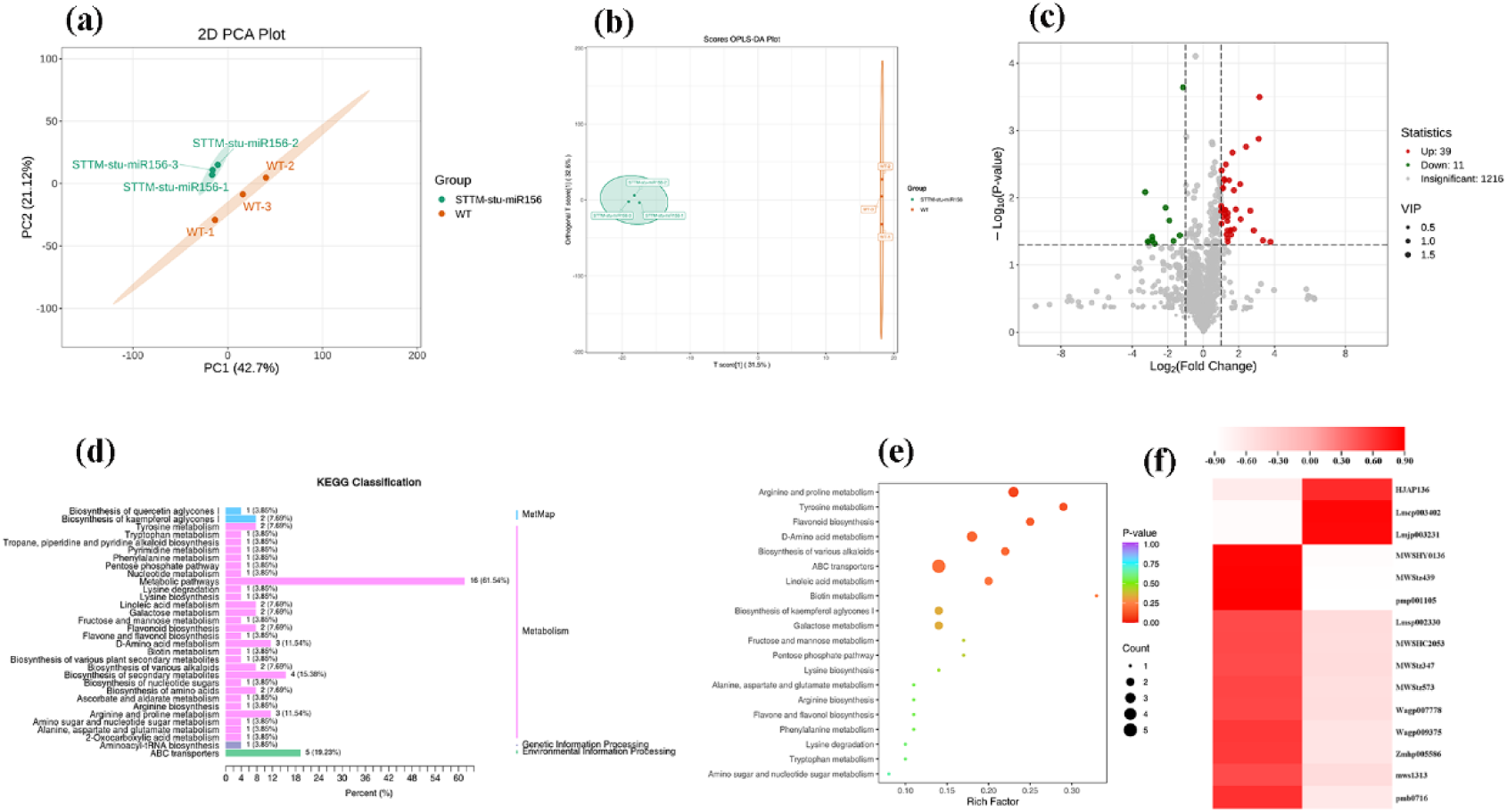
Metabolomic analysis of wild-type and transgenic potato plants. **a**: Principal component analysis; **b**: partial least squares discriminant analysis; **c**: Volcanic map of differential metabolites; Note: Green points represent down-regulated differential metabolites, red points represent up-regulated differential metabolites, and gray points represent detected but not significantly different metabolites; **d**: KEGG pathway annotation of DEMs; **e**: KEGG pathways; Note: The abscissa represents the Rich Factor corresponding to each pathway, the ordinate is the name of the pathway, the color of the point reflects the P-value size, and the redder the more significant the enrichment. The size of the point represents the number of differential metabolites enriched; **f**: The heat map of 15 DEGs related to flavonoids in differential metabolites showed that the deeper the red, the higher the degree of up-regulation.

To delve deeper into the molecular mechanisms that regulate the high levels of anthocyanins in transgenic potato plants, we individually assessed the expression levels of structural genes associated with anthocyanin across five different pathways, namely Phenylpropanoid biosynthesis (Ko00940), Flavonoid biosynthesis (Ko00941), Anthocyanin biosynthesis (Ko00942), Flavone and flavonol biosynthesis (Ko00944), and Stilbenoid, diarylheptanoid and gingerol biosynthesis (Ko00945) of STTM-miR156a and OE-miR156a (Table S3). The transcript abundance of Flavonoid 3’-monooxygenase (*UFGT*) was increased by 11.5 folds in OE-miR156a transgenic potato tubers. In contrast, the transcript abundances of Phenylalanine ammonia-lyase (*PAL*), 4-coumaric acid-CoA ligase (*4CL*), Chalcone synthase (*CHS*), Chalcone isomerase-like protein 2 (*CHI*), Naringenin,2-oxoglutarate 3-dioxygenase (*F3H*) and Dihydro flavonol 4-reductase (*DFR*) in OE-miR156a transgenic potato tubers were decreased by 4.9 folds, 4.5 folds, 2.9 folds, 19.9 folds, 8.0 folds and12.3 folds (Figure 4h). Meanwhile, qRT-PCR was used to identify the relative expression of these 7 DEGs, and the results were consistent with those of RNA-seq (Figure 4f). Compared with the wild type, the transcriptional abundances of these 6 DEGs in STTM-miR156a transgenic potato tubers exhibited an upward trend, and only the transcriptional abundance of *HCT* was decreased (Figure 4g), which was in line with the qRT-PCR results. Additionally, it was found that 2 *PALs*, 1 *4CL*, 1 *CHS*, and 2 *UFGTs* were up-regulated in OE-miR156a, suggesting that the regulation of miR156a affected the expression of these structural genes, thereby participating in the metabolic synthesis of anthocyanins in potato (Figure 4h).

### STTM-miR156a reduced anthocyanin accumulation in transgenic potato

Given the conserved function of miR156a and its interaction with anthocyanin biosynthesis and metabolism in potatoes, we selected wild-type (WT) and STTM-miR156a samples for further metabolomics analysis. The first two principal components clearly distinguished six samples of WT and STTM-miR156a, accounting for 42.7% and 21.12% of the overall variability, respectively. The metabolites were categorized into two groups, revealing a distinct separation trend an<u>d</u> notable differences. The samples within each group exhibited good reproducibility (Figure 5a). The findings derived from the OPLS-DA analysis (Figure 5b) were employed for subsequent model testing in conjunction with the differential metabolite screening criteria established in this study. In total, 152 distinct differentially accumulated metabolites (DEMs) were observed in the transgenic plants, which included 62 upregulated and 90 downregulated DEMs (Figure 5c). Analysis through KEGG and MetMap of the notably distinct metabolites in STTM-miR156a transgenic potatoes indicated that these metabolites were predominantly enriched in various metabolic pathways, ABC transporters, and the biosynthesis of secondary metabolites, in contrast compared to the wild type (Figure 5d). Notably, the differentially accumulated metabolites in STTM-miR156a potato tubers were significantly enriched in pathways such as arginine and proline metabolism, tyrosine metabolism, flavonoid biosynthesis, D-amino acid metabolism, and the biosynthesis of various alkaloids, flavone, and flavonol (Figure 5e).

The 15 differentially expressed metabolites (DEMs) associated with STTM-miR156a potato tubers, which participate in the biosynthesis of flavonoids and anthocyanins, are primarily categorized into two distributions (Figure 5f). 6 DEMs were screened from the ko00941 flavonoid biosynthesis pathway, including Alcesefoliside (5.5 folds), 5-Demethoxynobiletin (6.4 folds), Isosinensetin (2.2 folds), Quercetin 3,5,7,3,4-pentamethyl ether (8.7 folds), Sinensetin (5,6,7,3’,4’-pentamethoxyflavone)* (75.3 folds) and Tricin-4’-O-glucoside-7-O-glucoside* (57.5 folds). 9 DEMs were screened from the ko00944 flavone and flavonol biosynthesis pathway, including Quercetin-3-O-(2’’-O-rhamnosyl)rutinoside-7-O-glucoside (0.4 folds), Tamarixetin-3-O-glucoside (0.1 folds), Patuletin-3-O-glucoside (0.1 folds), Kaempferol-3-O-glucoside (Astragalin)* (2.6 folds), Camelliaside A (2.4 folds), Kaempferol-3-O-neohesperidoside-7-O-glucoside* (3.1 folds), Gossypetin 3,5,7,8,3’-pentamethyl ether (65.5 folds), 3,5,6,7-Tetramethoxyflavone (55.7 folds) and Herbacetin-7-O-(3’’-glucosyl) rhamnoside, Rhodiosin (15.8 folds). Among the 15 DEMs, the flavonoid-related metabolites, namely Quercetin-3-O-(2"-O-rhamnosyl) rutinoside-7-O-glucoside, Tamarixetin-3-O -glucoside (Tamarixin) and Patuletin-3-O-glucoside, were significantly reduced in comparison with the wild type. The remaining 12 DEMs, however, were significantly increased, which confirmed our total flavonoid quantitative results (Figure 3d and Table S4). Three anthocyanin-related metabolites, Cyanidin-3-O-glucopyranoside, Petunidin-3-O-rutinoside-5-O-glucoside, and Malvidin-3-O- rutinoside -5-O-glucoside, were decreased than the wild type (Table S4).

22 DEMs associated with phenolic acids were screened. Among these, the majority exhibited a decrease in phenolic acid content (Figure S3), which may be linked to a reduction in anthocyanin content in STTM-miR156a. In summary, compared to the wild type, the flavonoid levels in STTM-miR156a transgenic potato were significantly increased, while the levels of anthocyanins and phenolic acids were decreased. It indicates that miR156 plays a role in the metabolism of anthocyanins, phenolic acids, and flavonoids.

### DEGs and DEMs affected by miR156a

The correlation enrichment heatmaps of the top 20 DEGs and 20 DEMs (Table S4), identified between the transcriptome and metabolome, along with their corresponding correlation coefficients (represented in various colors), are presented in Fig. 6a. The upregulation of 17 DEGs resulted in the downregulation of the accumulation of 2-Phenylacetamide*, 3’-Glucosyl-6,7-dihydroxy-N-methyl-benzyltetrahydroisoquino -line, Leu-Ala-Trp and Cis-L-3-hydroxyproline*. Conversely, the downregulation of WRKY DNA-binding transcription factor 70 (LOC102577893), hypothetical protein KY289_026347 (novel.351), and MADS-box protein SVP (STMADS11) contributed to an increase in the accumulation of 17 DEMs. Additionally, the correlation network between DEGs and DEMs within the flavonoid (Ko00941) and flavonoid and flavonol (Ko00944) biosynthesis pathways were illustrated in Figure 4h (Figure 6b). A strong correlation was observed between 18 DEMs and 17 DEGs. The DEGs and DEMs associated with the flavonoid biosynthesis pathway (Ko00941) and the flavonoid and flavonol biosynthesis pathway (Ko00944) were illustrated in the enrichment heatmap (Figure 6c). Notably, Shikimate O-hydroxycinnamoyltransferase (*HCT*) and *4CL* exhibited a negative correlation with the majority of flavonoid-related metabolites. The remaining 15 DEGs were found to be negatively correlated with Patuletin-3-O-glucoside, Tamarixetin-3-O-glucoside, Phloretin-2’-O-6"-O-rhamnoside glucoside and Quercetin-3-O- (2"-O-rhamnosyl) rutin-7-O-glucoside, while they exhibited a positive correlation with the other 14 flavonoid-related DEMs.

**Figure 6.**
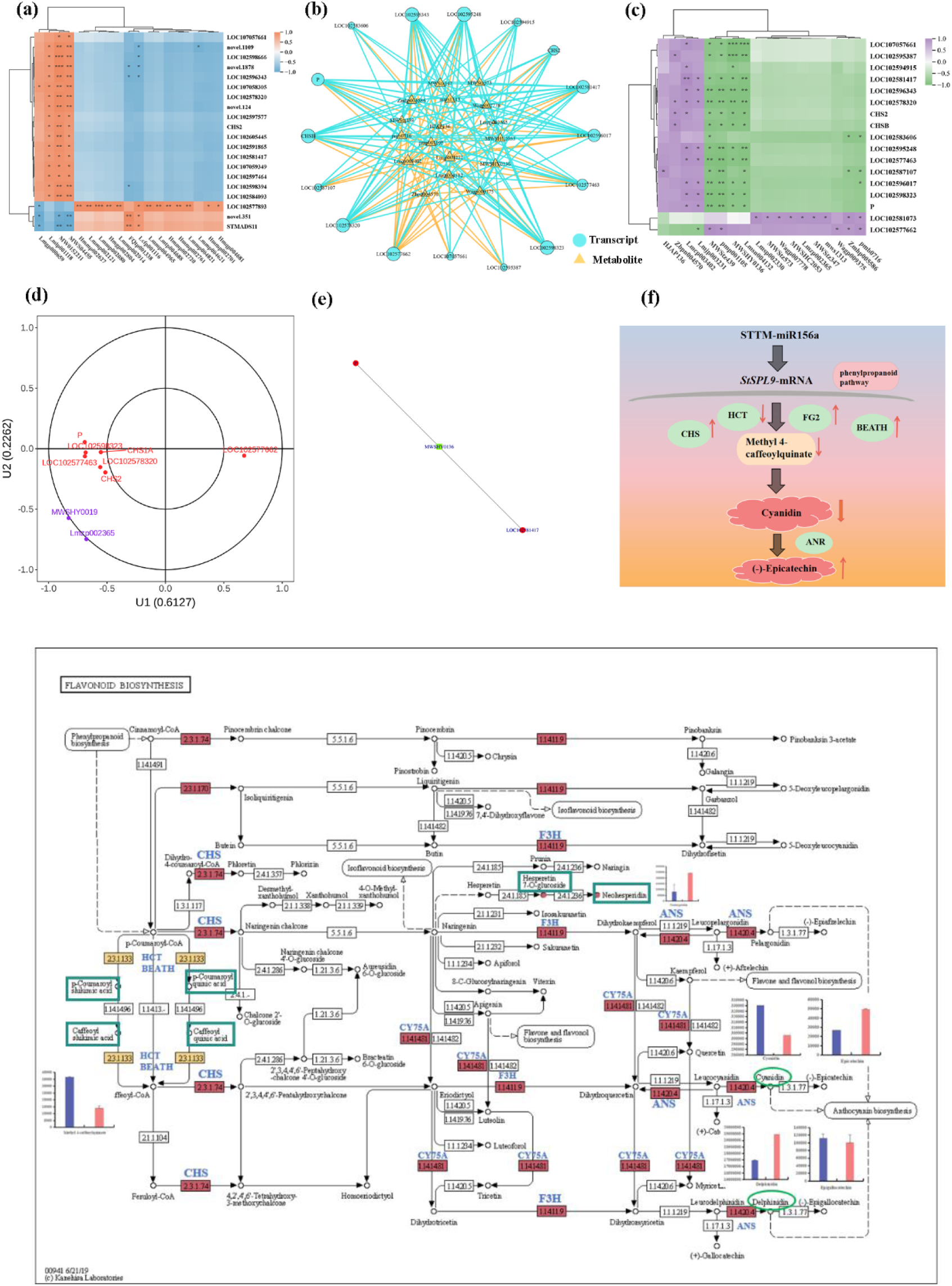
Joint metabolomic analysis of the transcriptome of STTM-miR156a transgenic potato. **a:** Heat maps of the first 20 enriched DEGs and DEMs before the STTM-miR156a transcriptome and metabolome; **b:** The network of DEGs and DEMs involved in flavonoid and anthocyanin biosynthesis pathways; **c:** DEGs and DEMs enrichment heat maps involved in flavonoid and anthocyanin biosynthesis pathways; **d:** CCA diagram of Ko00941 flavonoid pathway; Note: The horizontal and vertical coordinates represent the simple correlation coefficients between different omics substances and the typical variable U1 and the typical variable U2, respectively. Purple and red represent metabolites and genes, respectively. In the figure, persimmons are divided into four regions. In the same region, the farther the distance is, the stronger the correlation with typical variables is. The closer the distance to each other, the same correlation with the typical variables; **e:** The correlation network diagram of DEGs and DEMs with absolute value of Pearson correlation coefficient greater than 0.8 and pvalue less than 0.05 in Ko00944 flavonoid and flavonol pathway was selected; Note: The metabolites in the figure are marked with green squares, and the genes are marked with red circles. The solid line represents positive correlation, and the dotted line represents negative correlation; **f:** A model was constructed to explain how miR156a and its target genes regulate anthocyanin biosynthesis in potato tubers; **g:** Biosynthesis and metabolic pathways of flavonoids in STTM-miR156a potato tubers (non-transgenic potato as control group).

Through canonical correspondence analysis (*CCA*) and the construction of related networks, it was determined that flavonoid 3’,5’-hydroxylase (*CYP75A2*), leucoanthocyanidin dioxygenase (*ANS*), chalcone synthase (*CHS*), flavanone 3-hydroxylase (*F3H*), acetyl-CoA-benzyl alcohol acetyltransferase-like (*BEATH*) were involved the flavonoid biosynthesis pathway (Ko00941) and the flavonoid and flavonol pathway Ko00944. A total of 5 DEGs in anthocyanidin 3-O-glucosyltransferase (*FG2*) were shown to influence the accumulation of three differential metabolites, namely Hesperetin 7-O-glucoside, Hesperetin-7-O-neohesperidoside and Kaempferol 3-O-glucoside (Table S4), by up-regulating and down-regulating the expression of *HCT*. Additionally, network analysis revealed that *CYP75A2* and *FG2* co-regulated Kaempferol-3-O-glucoside (Astragalin). The accumulation of two colorless metabolites, Hesperetin 7-O-glucoside and Neohesperidin, was enhanced in STTM-miR156a, whereas the accumulation of Kaempferol 3-O-glucoside was inhibited (Figure 6d e).

Methyl 4-caffeoylquinic acid ester was a type of plant secondary metabolite (Table S4). Its synthesis involves a catalytic reaction mediated by various enzymes and was a key component of the phenylpropanoid metabolic pathway. Notably, it was observed that *StSPL9*, *HCT*, *BEATH*, and *FG2* collectively reduced the content of methyl 4-caffeoylquinic acid ester. The low expression of *HCT* combined with the high expression of *BEATH* resulted in decreased accumulation of methyl 4-caffeoylquinate, which may contribute to reduced cyanidin levels due to the actions of *CHS*, *F3H*, and *ANS* or decreased delphinidin might stem from the collective influence of *CHS*, *CY75A*, *F3H* and *ANS* (Figure 6g). In summary, the accumulation of methyl 4-caffeoylquinate was likely diminished due to the high expression of *BEATH* and *FG2*, in conjunction with the low expression of *HCT* (Figure 6f).

## Discussion

Colored potato varieties, characterized by their anthocyanin-rich skins and tubers, provide protection to the human body against oxidants, free radicals, and LDL cholesterol. The antioxidant activity of potato tubers primarily arises from flavonoids, anthocyanins, and phenolic acids (Keutgen et al, 2019). Anthocyanins have been extensively studied in various plant species, and the mechanisms underlying their accumulation have been elucidated (Zhou et al, 2022). Guo demonstrated that the overexpression of miR156b in grape cells increased anthocyanin content, while STTM-miR156b produced the opposite phenotype. In this study, STTM-miR156a and OE-*StSPL9* transgenic plants exhibited lower anthocyanin content, whereas OE-miR156a potato leaves and tubers displayed the opposite phenotype (Figure 3) (Wang et al, 2020), consistent with earlier findings. These investigations suggest that miR156 possesses both conserved and species-specific traits among various species.

A previous study demonstrated that miR156 positively regulates anthocyanin accumulation in poplar (Zhou et al, 2022; Gou et al, 2024). miR156a is a highly conserved miRNA across evolutionary contexts. However, the precise mechanisms by which miR156a contributes to anthocyanin accumulation in potatoes remain poorly understood. This study, presents several lines of evidence indicating that miR156a was a critical gene that positively regulates anthocyanin accumulation in potatoes by repressing its target gene, *StSPL9*. Firstly, our results showed that overexpression and silencing of miR156a resulted in increased and decreased anthocyanin content of potato tubers, respectively (Figure 3d), which was in agreement with the results reported by Wang in poplar (Wang et al, 2020). Second, miR156a primarily influences anthocyanin accumulation through the regulation of its target gene *StSPL9*. After overexpression or silencing of miR156a, the expression of *StSPL9* was down-regulated or up-regulated respectively (Figure 3b and c). Furthermore, overexpression of *StSPL9* significantly reduced anthocyanin content in potato tubers (Figure 3h). Luo demonstrated that the expression level of *StSPL9* was down-regulated in overexpressing miR156 transgenic plants (Lou et al, 2024). Thus, *StSPL9* is identified as the primary target gene of miR156a in the context of anthocyanin accumulation in potato tubers. Given that several other target genes, including *StSPL1*, *StSPL3*, *StSPL6*, *StSPL7*, *StSPL8*, *StSPL9,* and *StSPL12* were also influenced by the overexpression of miR156a (Figure 4e), it cannot be rule out that other *StSPL* members may act synergistically in the process of anthocyanin accumulation in potato tubers (Li et al, 2024; Li et al, 2020). It has been established that the target gene of miR156 is *SPL* in *Arabidopsis*, rice, and other plants species (Gandikota et al, 2007; Xie et al, 2006; Liu et al, 2024). For instance, in apple fruit and callus, the overexpression of *MdSPL2-like* and *MdSPL33* promotes anthocyanin accumulation, while silencing these genes yields opposite results (Yang et al, 2024). Similarly, in *Salvia miltiorrhiza*, overexpression of *SmSPL7* also enhances anthocyanin accumulation (Chen et al, 2021). In summary, the miR156/*SPL9* model plays a crucial role in regulating anthocyanin biosynthesis in potato tubers. The expression levels of miR156a and *StSPL9* correlate with the changes in potato anthocyanin-related gene expression, thereby influencing the concentrations of related compounds in potato tubers. This provides a significant reference value for the molecular breeding of colored potatoes.

More and more scholars have proved that miR156 is related to regulating anthocyanin biosynthesis (Lin et al, 2024; Liu et al, 2017). The MBW complex positively regulates the expression of structural genes by binding to cis-acting elements in gene promoter regions (such as *DFR*, *LODX* /*ANS*, *UFGT*, etc.) and then promotes the biosynthesis of anthocyanins in plant species (Ni et al, 2020; Xu et al, 2015). *PAL* is the first structural gene involved in the biosynthesis of anthocyanins, with Wang (Wang, 2022) demonstrating that *PAL* activity can enhance the formation of anthocyanins and flavonoids. In this study, the expression of *StPAL* was up-regulated in STTM-miR156a, while it was down-regulated in OE-miR156a. Ma (Ma et al, 2023) revealed in research on the function and structure of *St4CL* in apples that the expression level of *St4CL* is positively correlated with the accumulation of anthocyanins. In this study, *St4CL* displayed elevated expression levels in both STTM-miR156a and OE-miR156a. Shi (Shi et al, 2021) discovered that *CHI*, *CHS*, and *FLS* interact to modulate the biosynthesis of flavonols. In STTM-miR156a, both *StCHI* and *StCHS* were up-regulated, whereas an opposite trend was observed in OE-miR156a. This indicates that a greater quantity of flavonoid compounds may be synthesized in STTM-miR156a (Figure 4h).

*F3H* is a relay gene involved in anthocyanin biosynthesis, and its high expression can result in reddish-brown petals in dahlias (Gutiérrez-Albanchez et al, 2020). The high expression of *StF3H* in STTM-miR156a could play a role in the synthesis of anthocyanins in potato tubers, especially when contrasted with the decreased expression observed in OE-miR156a. Regarding *DFR*, the up-regulation of *DFR* upstream facilitates the movement of substrates towards the flavonol pathway (Li et al, 2022). In this study, *StDFR* was found to be up-regulated in STTM-miR156a, while it was down-regulated in OE-miR156a. Yan reported that the up-regulation of *ANS* increased anthocyanin content in poplar (Yan et al, 2022). Notably, *StANS* emerged as the key gene influencing anthocyanin formation, thereby determining the coloration of potato tubers. In our research, *StANS* was highly expressed in both overexpressed and silenced transgenic potato plants, further catalyzing the formation of anthocyanins from substrates in both silenced and overexpressed plants. The phenotypic results indicated that the silenced transgenic potato tubers exhibited a lighter lavender color. We discovered that *StANS* upregulates expression in the cyanidin synthesis pathway, resulting in an increased content of Cyanidin-3-O-glucosylrutinoside metabolites. Most of the Cyanidin-3-O-glucosylrutinoside is subsequently converted to (-)-Epicatechin through the action of *StANR* (LOC102600995), ultimately leading to a reduction in anthocyanin content (Figure 6g).

Gene expression is closely linked to the synthesis of metabolites. The distinct characteristics of plants can be attributed to variations in the gene regulation of these metabolites. Upon the inhibition of miR156a expression, there was an upregulation of *CYP75A*, *ANS*, *CHS*, and *F3H*, while the expression of *HCT* was down-regulated. This resulted in the promotion of the accumulation of two flavone metabolites, namely Hesperetin 7-O-glucoside and Neohesperidin, while the accumulation of Kaempferol 3-O-glucoside was inhibited. Through the construction of relevant networks, it was revealed that *CYP75A2* and *FG2* co-regulate Kaempferol-3-O-glucoside (Astragalin).

Down-regulated expression of *CYP75A2* may affect anthocyanin accumulation in eggplant (*Solanum melongena* L). This finding suggests that *CYP75A2* and *FG2* may be crucial regulatory factors governing anthocyanin accumulation in potatoes (Figure 6d and e), which could potentially explain the difference in potato tuber color.

According to existing literature, compelling evidence suggests that pigmented tissues contain nearly twice the quantity of phenolic acids compared to non-pigmented tissues (Navarre et al, 2013). Phenolic acids, with chlorogenic acid (*CGA*) as the predominant constituent, represent the principal class of compounds, followed by anthocyanins and flavan-3-ols (Valiñas et al, 2017), in *Solanum tuberosum subsp. Andigena*, *HCT* is involved in producing of chlorogenic acids (Valiñas et al, 2017). Furthermore, in *Ligularia fischeri*, a favorable regulatory relationship exists between methyl 4-caffeoylquinate and chlorogenic acids (Park et al, 2023). Research by Qian has shown that the expression of *HCT* in yellow-tubers potatoes is significantly lower than in colored potatoes, and the accumulation patterns of chlorogenic acid are highly correlated with those of 4-caffeoylquinic acid (Qian et al, 2017). A decrease in *HCT* reduces the content of caffeoylquinic acids (CQAs) (Park et al, 2023). This study found that the expression of hydroxycinnamoyl transferase (*StHCT*) was decreased in STTM-miR156a, while no such change was observed in OE-miR156a. Notably, methyl 4-caffeoylquinate is a type of plant secondary metabolite synthesized by plants through their endogenous biosynthetic pathways. This synthesis process involves catalytic reactions mediated by multiple enzymes and is a crucial component of the phenylpropanoid metabolic pathway. The findings indicate that a reduction in *StHCT* expression leads to lower levels of methyl 4-caffeoylquinate, which consequently causes an increase in the downstream accumulation of delphinidin and cyanidin (Figure 6g and Figure S3). Thus, it is quite likely that the *StHCT* gene is crucial and central to the biosynthesis of CQAs in potatoes.

## Conclusion

The findings indicated that miR156a promoted the accumulation of total anthocyanins in potato tubers, while *StSPL9* inhibited this accumulation. In STTM-miR156a and OE-miR56a, the differential expression of anthocyanin biosynthetic structural genes and regulatory genes resulted in differences of metabolite, thereby causing changes in the tuber color of these transgenic potatoes. In STTM-miR156a transgenic potatoes tubers, the contents of most flavonoid metabolites have increased, whereas the contents of most phenolic acids have decreased. Phenolic acids contribute to the stability of anthocyanins enhance color intensity (Valiñas et al, 2017). This may be an additional factor contributing to the lighter color of potato tubers.

## Material and moehods

### Plant materials and the growth conditionals

In this experiment, sterile tissue culture seedlings of *Nicotiana benthamiana* and purple potato (*Solanum tuberosum* L.) variety ‘Huasong 66’ were used as experimental materials for genetic transformation of wild-type materials (WT). The potato tissue culture seedlings were provided by Inner Mongolia Huasong Agricultural Science and Technology Co, Ltd. The test plants were propagated on MS medium and cultured at 25±2°C for 16/8h under light/dark cycle. The relative humidity (RH) was 60 %. The culture for about 3-4 weeks can be used for subsequent experiments. The experimental materials for transcriptomics and metabolomics were planted in the greenhouse of the Agricultural College of Inner Mongolia Agricultural University (Hohhot, China, the same culture conditions as the tissue culture room) in July 2023. Scientific management was carried out during the planting period to ensure the normal growth of transgenic potato plants. The samples were collected at the maturity stage of potato tubers, and each sample was repeated three times. After harvesting the mature tubers of potatoes, they were immediately frozen in liquid nitrogen after sampling and then stored in a refrigerator at -80 ° C for subsequent experiments.

### Vector construction and transformation of miR156 and its target gene *StSPL9*

The base sequence of miR156a was obtained by referring to the sequencing of the Small RNA degradation group in our laboratory (Wu et al, 2022). The specific (F/R) primers were designed to clone the miR156a gene from the cDNA of ‘the Huasong 66’ tuber. The miR156a was ligated into the silencing vector pBWA (V) KS-8601 and the overexpression vector pCAMBIA1302 by homologous recombination and golden gate seamless cloning to verify whether the actual fragment size of the recombinant plasmid was consistent with the theory. The recombinant plasmids pBWA (V) KS-miR156a and pCAMBIA1302-miR156a were sequenced (Sangon, Shanghai, China). After sequencing, T4 DNA ligase was ligated, and then the ligation products pBWA (V) KS-miR156a and pCAMBIA1302-miR156a were transformed into *Escherichia coli* TOP10 competent cells. Positive single colonies were screened on LB solid medium containing 50μg/mL kanamycin, and the single colonies with good growth were identified by PCR and sent to sequencing (Sangon, Shanghai, China). The silencing vector pBWA (V) KS-miR156a and pCAMBIA1302-miR156a containing the 35S promoter were correctly sequenced and introduced into the *Agrobacterium* strain LBA4404. The pCAMBIA1302-*StSPL9* expression vector was constructed by double enzyme digestion. In this study, Agrobacterium-mediated genetic transformation of potatoes was carried out. The sterile tissue culture seedlings of potato that have been cultured for 3-4 weeks were selected, and the stem segments were cut and cultured until the regenerated seedlings resistant to hygromycin and kanamycin were differentiated. After acclimation, the rooted seedlings were transplanted into nutrient soil. Fast Pure Plant DNA Isolation Mini Kit-BOX2 kit was used to extract DNA from transgenic potato tubers, and STTM-35seq-F/NOS seq-R and pCAMBIA1302 miR156-F/R primers were used to identify positive plants.

### Verification of the interaction between miR156a and its target gene StSPL9

The recombinant positive vectors pCAMBIA1302-StSPL9-mGFP5 and pCAMBIA1302-miR156a-mGFP5 were activated and resuspended. The pCAMBIA1302-mGFP5, pCAMBIA1302-StSPL9-mGFP5, pCAMBIA1302-miR156a-mGFP5, the equal volume mixed suspension of pCAMBIA1302-StSPL9-mGFP5 and pCAMBIA1302-miR156-mGFP5 was injected into *Nicotiana benthamiana* leaves cultured at room temperature for 2h, respectively. After 1d of dark and light culture after injection, the fluorescence of leaves was observed under a fluorescence confocal microscope, and RNA was extracted. qRT-PCR detected the relative expression of mGFP5.

### Total RNA extraction and reverse transcription and real-time fluorescence quantitative PCR

Extraction of potato tuber RNA by Trizol method: TRIzol ® Reagent was added to the powder ground from potato tubers. After adding chloroform to stand, the upper layer containing RNA was extracted. RNA was further precipitated with isopropanol, and impurities such as DNA or protein were washed to obtain total RNA. The quality of total RNA was tested. The quality control range was: OD260 / 280, which was between 1.8-2.2 and 28S: 18S ≥ 1.5. The instrument used for qRT-PCR was (Thermo Fisher Scientific, Quant StudioTM5). miR156a used potato U6 as an internal reference gene and the miRcute enhanced miRNA cDNA first-strand synthesis kit (TIANGEN LOTY2109, Beijing, China) to synthesize cDNA by reverse transcription. The expression of miR156a was detected by miRcute enhanced miRNA fluorescence quantitative detection kit (TANGEN FP411, Beijing, China). *StSPL9* used potato Actin as the internal reference gene (Supplementary Table.S2), and the expression of the target gene was detected using the MonAmpTM SYBR ® Green qPCR Mix kit from Monad Biotechnology Co, Ltd. The PCR reaction procedure included initial denaturation at 95 °C for 10s, followed by 40 cycles, and annealing at 60°C for 30s. The dissolution curve was collected using the instrument’s default procedure.

The calculation formula of qRT-PCR detection of transgenic potato was RQ (relative expression) = 2^-ΔΔCT^ (Livak et al., 2001). Comparing the gene expression levels of wild-type and transgenic plants, the detection method of relative expression of miR156a and its target gene *StSPL9* was calculated according to qRT-PCR detection reference 2^-ΔCT^ (Dheda et al., 2004). To minimize errors and enhance the reliability of results, we conducted three biological and three technical replicates. For the one-way analysis of variance, the Duncan multiple range test was executed using SPSS Statistics 26.0 software, and a significance level of P < 0.05 was applied.

### Determination of phenotypic and physiological indexes of wild-type and transgenic potato tubers

We determined the total anthocyanin and flavonoid contents of the samples analyzed by transcriptomics and metabolomics. Anthocyanin content was detected using a plant anthocyanin content detection kit (Suolaibao BC1385, Beijing, China). Anthocyanins were extracted from 0.1 g of fresh potato tubers and leaves. A microplate reader (BioTek, EPOCH, USA) was employed to measure the absorbance of the extract at wavelengths of 530 and 700 nm. In addition, the flavonoid content in the tubers from both wild-type and transgenic potatoes was detected by the plant flavonoid content detection kit (Suolaibao BC1335 Beijing, China). All procedures were carried out strictly following the instructions provided with the kit. The absorbance for flavonoid quantification was also measured at 470 nm using the microplate reader.

### Transgenic potato transcriptome sequencing and analysis

Transcriptome analysis was performed on 3 wild-type potato tubers (WT), 3 STTM-miR156a, and 3 OE-miR156a transgenic potato tubers. RNAprep Pure Plant Total RNA Extraction Kit (Trelief ® RNAprep Pure Plant Plus Kit, DP441; tianGen; Beijing, China) to extract total RNA from potato tubers. The concentration of RNA was measured by Qubit 4.0 fluorescent agent/MD microplate reader, the integrity of RNA was measured by Qsep400 biological analyzer, and the better-quality RNA was reversely transcribed into cDNA. The Illumina Hiseq (Illumina NovaSeq 6000) platform (Metware Biotechnology Ltd; Wuhan, China) for sequencing. The clean reads used in the subsequent analysis were obtained by Fastp (Chen et al., 2018) for raw data filtering, sequencing error rate checking, and GC content distribution checking.

For the samples with three biological replicates, DESeq2 (Buchfink et al., 2015; Zheng et al., 2016) was used to analyze the differential expression between the sample groups, and the Benjamini-Hochberg method was used to perform multiple hypothesis test correction on the hypothesis test probability (P value) to obtain the false discovery rate FDR (False Discovery Rat). Differential genes were identified based on the criteria of | log2Fold Change | ≥ 1 and FDR < 0.05. Annotation of these differential genes was carried out using the KEGG database (https://www.genome.jp/kegg) and the GO database (http://www.geneontology.org). Enrichment analysis was conducted using the hypergeometric test. In the case of KEGG, pathways served as the units of analysis, while for GO, the analysis relied on GO entries.

### Metabolomics sequencing and analysis of transgenic potato

Three wild-type potato tubers and three transgenic tubers expressing STTM-miR156a were chosen for metabolomics evaluation. The samples underwent freeze-drying utilizing a freeze-dryer (Scientz-100F), were pulverized with a grinder (MM400, Retsch) operating at 30HZ, and centrifuged in a centrifuge (5424R, Hamburg, Germany). The resulting supernatant was gathered and filtered through a microporous membrane with a pore size of 0.22μm for further UPLC-MS/MS analysis. Ultra-performance liquid chromatography (UPLC) (ExionLC TM AD, https://sciex.com.cn/) was employed, combined with tandem mass spectrometry (MS/MS). The QQQ scan operated in MRM mode, with the collision gas (nitrogen) set to a medium level. The multiple reaction monitoring (MRM) mode, acquired via triple quadrupole mass spectrometry and informed by the malware database (MWDB), facilitated the detection of metabolites within the analyzed samples. To ensure the reliability of both qualitative and quantitative assessments of the metabolites, the mass spectrum peaks from each identified metabolite across the six samples were integrally adjusted. Throughout the process of instrumental analysis, quality control (QC) samples were incorporated to evaluate the technical reproducibility of metabolite extraction and detection. To assess the overall metabolic variance among all identified metabolites, strains, and various biological replicates, principal component analysis (PCA) and orthogonal partial least squares discriminant analysis (OPLS-DA) were applied.

### Joint analysis of transcriptome and metabolome

The reasons for the formation of potato anthocyanins were analyzed from the results of overall differential metabolites and transcriptome differential genes. A pearson correlation analysis was conducted on the correlation coefficients of genes and metabolites identified in each differential group. Results with absolute correlation coefficient values exceeding 0.8 and p-value under 0.05 were chosen for further analysis, including the correlation coefficient clustering heat map and correlation network diagram. To integrate and evaluate the overall correlation between metabolomics and transcriptomics data, the two-way orthogonal partial least squares (O2PLS) model was employed.

### Statistical analysis

In this study, three wild-types and three STTM-miR156a -10, -12, -13, three OE-miR156a -7, -10, -12, and three OE-*StSPL9*-1, -7, -14 transgenic potato tubers were analyzed using Microsoft Excel 2019 (Microsoft, Washington, DC, United States) and SPSS 23 (SPSS Inc., Chicago, IL, United States) for statistical analysis. To assess the significance among the samples, a one-way analysis of variance (ANOVA) was performed, with a significance threshold set at P < 0.05. To minimize potential errors and enhance the reliability of our findings, we conducted three biological replicates and three technical replicates throughout the experiment. Additionally, standard deviation (STDEV) was employed to measure and adjust for significant disparities among the data sets.

## Acknowledgements

This study was supported by grants from Natural Science Foundation of Inner Mongolia Autonomous Region of China (2022JQ06), Breeding Joint Research Project of Inner Mongolia Autonomous Region of China (YZ2023006), Special Funds for Basic Scientific Research Operating Expenses of Inner Mongolia Agricultural University (BR22-11-16) and Postgraduate Scientific Research Innovation Project of Inner Mongolia Autonomous Region (S20231102Z). Funders had no role in the design of the study and collection, analysis, and interpretation of data and in writing of the manuscript.

## Supplementary Data

**supplementary Figure S1.** Baining and identifying of StumiR156a transgenic potato plants.

**Supplementary Figure S2.** KEGG clustering map and differentially expressed transcription factors in transgenic tubers.

**Supplementary Figure S3.** Enrichment heatmap of phenolic-acid-related differential metabolites.

**Supplementary Table S1.** Primers used for cloning and vector construction detection.

**Supplementary Table S2.** Primers used in this study for quantitative analysis.

**Supplementary Table S3.** Gene IDs and information of SPLs within the transcriptome datasets of STTM- miR156a and OE-miR156a.

**Supplementary Table S4.**Names and Classifications of Metabolites in STTM-miR156a.

## Author contributions

All authors participated in the study and agreed with the content of this manuscript. Y.H.M. conceived and designed the research, N.L., P.J.W., Y.M. and J.W, carried out experiment, N.L. wrote the manuscript, N.L., X.J.W., H.S.N. and E.Z.Y. participated in experiments, N.L., Z.C.Z, D.W. and X.R. assisted with data analysis.

## Data availability statement

Genes sequence and RNA-Seq data are available from NCBI BioProject ID PRJNA1191835 (http://www.ncbi.nlm.nih.gov/bioproject/1191835). Additional relevant data can be found in this manuscript or in the supplementary material.

